# Abundance estimation from genetic mark-recapture data when not all sites are sampled: an example with the bowhead whale

**DOI:** 10.1101/549394

**Authors:** Timothy R. Frasier, Stephen D. Petersen, Lianne Postma, Lucy Johnson, Mads Peter Heide-Jørgensen, Steven H. Ferguson

**Affiliations:** Department of Biology, Saint Mary’s University, 923 Robie St., Halifax, NS, B3H 3C3 Canada; Conservation and Research Department, Assiniboine Park Zoo, 2595 Roblin Blvd., Winnipeg Manitoba, R3R 0B8, Canada; Fisheries and Oceans Canada, 501 University Crescent, Winnipeg, Manitoba, R3T 2N6, Canada; Greenland Institute of Natural Resources, Box 570, 3900 Nuuk, Greenland

**Author notes:** **Corresponding Author:** Timothy R. Frasier, Saint Mary’s University, Department of Biology, 923 Robie Street, Halifax, NS B3H 3C3, Canada.

**Keywords:** abundance estimation, mark-recapture, genetic identification, bowhead whale, whale, genetic mark-recapture

## Abstract

Estimating abundance is one of the most fundamental and important aspects of population biology, with major implications on how the status of a population is perceived and thus on conservation and management efforts. Although typically based on one of two methods (distance sampling or mark-recapture), there are many individual identification methods that can be used for mark-recapture purposes. In recent years, the use of genetic data for individual identification and abundance estimation through mark-recapture analyses have increased, and in some situations such genetic identifications are more efficient than their field-based counterparts for population monitoring. One issue with mark-recapture analyses, regardless of which method of individual identification is used, is that the study area must provide adequate opportunities for “capturing” all individuals within a population. However, many populations are unevenly and widely distributed, making it unfeasible to adequately sample all necessary areas. Here we develop an analytical technique that accounts for unsampled locations, and provides a means to infer “missing” individuals from unsampled locations, and therefore obtain more accurate abundance estimates when it is not possible to sample all sites. This method is validated using simulations and is used to estimate abundance of the Eastern Canada-West Greenland (EC-WG) bowhead whale population. Based on these analyses, the estimated size of this population is 11,747 individuals, with a 95% highest density interval of 8,169-20,043.

## 1. Introduction

Mark-recapture analyses have become an invaluable tool not only as a means for abundance estimation, but also for obtaining a wealth of other information on population dynamics, such as population growth rates, life history, and survival probabilities (Kraus et al. 2001; Williams et al. 2002; Amstrup et al. 2005). Traditionally, mark-recapture methods involved the actual capture, marking, and recapture of individuals (Rice and Harder 1977; Hestbeck et al. 1991; Pradel et al. 1997), but their use quickly expanded to species where individuals could be identified based on natural markings, and thus “capturing” in these cases meant taking a picture that could be used for identification (Wilson et al. 1999; Matthews et al. 2001; Langtimm et al. 2004; Bradshaw et al. 2007). More recently, the increasing ease with which individual-specific genetic profiles can be obtained from small sources of DNA has made it possible (and sometimes more efficient) to use genetic data as the primary means of individual identification, and thus the basis for mark-recapture analyses (Mowat and Strobeck 2000; Miller et al. 2005; Petit and Valiere 2006).

One important requirement in mark-recapture studies, regardless of which method of identification is used, is ensuring that the study area provides adequate opportunities for “capturing” all individuals within the study population. This requirement may be easily addressed for populations with small ranges that can be surveyed in their entirety, or where all individuals utilize—or pass through—a single area. However, many populations are not so amenable, with individuals exhibiting substantial variation in site fidelity and habitat use patterns. Failure to account for this variation can bias abundance estimates either upwards or downwards. The former can occur if marked individuals move out of the study area and thus become unavailable for recapture. The latter can occur if the study area is only used by a portion of the population, and thus estimates are only representative of some individuals rather than the entire population.

For species with wide distributions and substantial heterogeneity in habitat use patterns it may not be possible to obtain adequate coverage of all areas. In such cases abundance estimation requires appropriate corrections during the analytical stages of the study. In 2005, Durban et al. (2005) made substantial progress in this area by recognizing that analytically accounting for missed sites is essentially a regression problem, where data from sampled areas can be used to make predictions on the number of individuals not seen in any surveyed area. As an example, suppose a study population of organisms where some, but not all, individuals can be identified based on natural markings. Suppose the number of identifiable individuals has been counted at each of three sites, but other unstudied sites exist. To estimate the total size of the identifiable (marked) population, Durban et al. (2005) first organized the sighting histories of identifiable individuals based on whether or not they were observed in each of the three surveyed areas (see **Table 1**). The last row of such a table represents the identifiable individuals not seen in *any* location, and can be inferred from the available data. In this case a regression model is fit with the first seven rows of data, and the coefficient estimates from this process are then used to predict the value for the last row. In their application of this approach to a study of bottlenose dolphins *(Tursiops truncatus)* around Scotland, Durban et al. (2005) found that their approach performed quite well and resulted in a similar estimate to that obtained through another study based on a more rigorous and standardized survey design.

**Table 1.** A table showing the data layout as needed to infer the number of marked individuals in unsampled sites, as shown in Durban et al. (2005). Each row represents individuals with a particular capture history. For example, row one represents individuals seen in all three areas. Row two represents individuals seen in areas #1 and #2, but not area #3. Row three represents individuals seen in area #1 and area #3, but not area #2, and so on. The last row represents identifiable individuals not seen in any of the surveyed areas, and the number of individuals with this sighting history is to be inferred.

| Marked Individuals | Area 1 | Area 2 | Area 3 |
| --- | --- | --- | --- |
| 1 | 1 | 1 | 1 |
| 16 | 1 | 1 | 0 |
| 3 | 1 | 0 | 1 |
| 35 | 1 | 0 | 0 |
| 2 | 0 | 1 | 1 |
| 2 | 0 | 1 | 0 |
| 16 | 0 | 0 | 1 |
| ? | 0 | 0 | 0 |

The eastern Canada-western Greenland (EC-WG) bowhead whale *(Balaena mysticetus*, **Fig. 1**) population epitomizes the confounding factors associated with abundance estimation, using either distance sampling or mark-recapture methods. First, it is an arctic-adapted population with a wide distribution often associated with the ice edge (**Fig. 2**) (Reeves et al. 1983; Moore and Reeves 1993; Finley 2001; Ferguson et al. 2010). The necessary logistics and monetary requirements make extensive surveys in the high arctic challenging. Second, the population is highly segregated based on sex, age class, and perhaps other factors (Reeves et al. 1983; Finley 1990; Moore and Reeves 1993; Heide-Jørgensen et al. 2010). Indeed, the segregation is so great that, until recently, multiple populations (and subsequent management stocks) were thought to exist (Rugh et al. 2003). It was only when satellite tagging data recently showed movement of whales across previously assumed population boundaries, and such movements were also identified based on genetic mark-recapture analyses, that the presence of only one population was recognized (Heide-Jørgensen et al. 2003, 2006; Wiig et al. 2011). These confounding factors have led to very imprecise abundance estimates, as well as large differences in estimates based on data from different locations and/or different analytical approaches, as summarized in Koski et al. (2013).

**Figure 1.**
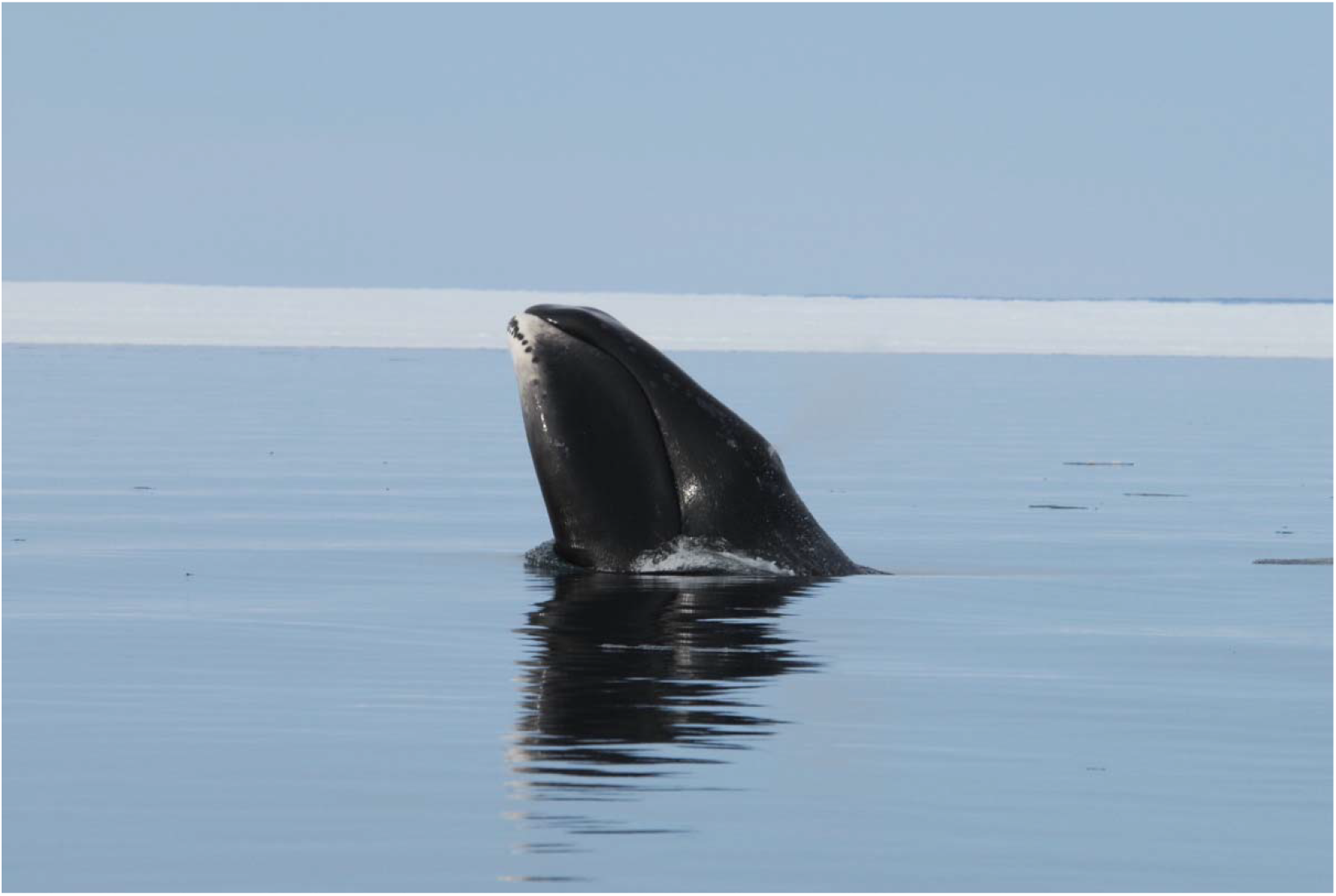
Bowhead whale *(Balaena mysticetus)*. Photograph by Jeff Higdon.

**Figure 2.**
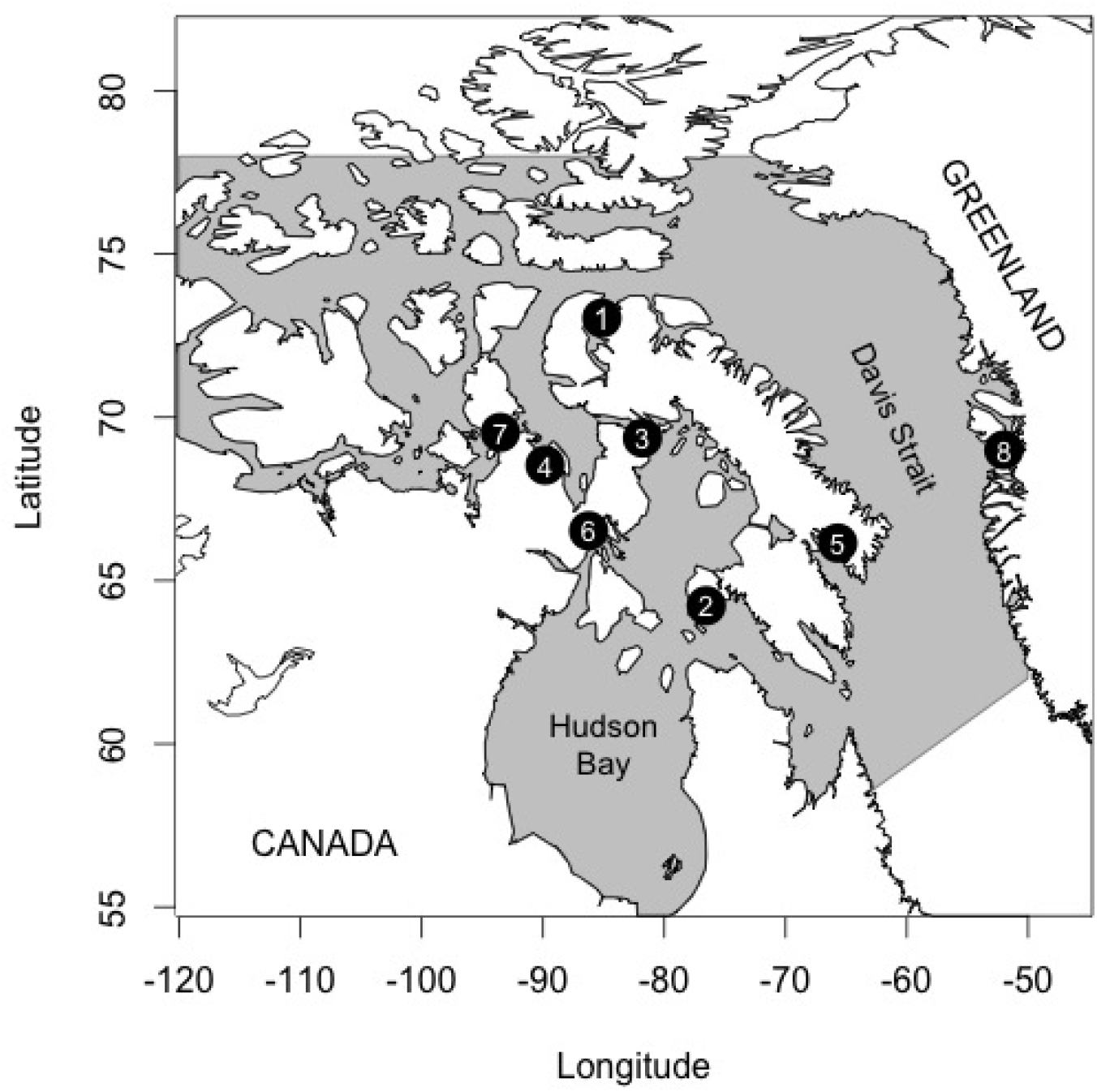
Distribution and sampling locations of the eastern Arctic bowhead population. Sampling locations are numbered as follows: 1 = Arctic Bay, 2 = Cape Dorset, 3 = Igloolik, 4 = Kugaaruk, 5 = Pangnirtung, 6 = Repulse Bay, 7 = Taloyoak, and 8 = Disko Bay.

Here, we extend the approach of Durban et al. (2005) in an important way. Specifically, The model of Durban et al. (2005) was designed to estimate the number of missing *marked* individuals in a population. However, our goal was to estimate the *total number* of missing individuals (marked and unmarked), and combine this information with data from sampled sites to result in one estimate of total abundance for the population of interest. If successful, this approach could prove useful to a broad range of population studies. We then applied this method to genetic mark-recapture data for the eastern Arctic-western Greenland bowhead whale population. Genetic mark-recapture methods may be more efficient than photograph-based identifications for this population because samples can be collected in collaboration with Arctic communities, eliminating the need for extensive and expensive aerial surveys while also fostering closer collaborations and relationships with Arctic communities.

## 2. Methods

### 2.1 Sampling

Intensive sampling effort has taken place out of the communities of Igloolik and Pangnirtung in Canada, and in Disko Bay, Greenland (hereafter “Greenland”) (**Fig. 2**, **Table S1**). Additionally, more opportunistic sample collection has occurred in Repulse Bay and Arctic Bay, as well as from 11 subsistence hunts by Canadian Inuit (Taloyoak, Kugaaruk, Cape Dorset, Pangnirtung, Repulse Bay, Igloolik) (**Fig. 2**, **Table S1**). In total, 1,177 samples have been collected across 19 years.

In each location, small tissue (skin and blubber) samples were obtained from free-swimming whales using a 150-lb draw-weight crossbow (Excalibur Vixen) with bolts and tips modified specifically for this purpose (from Ceta-DART, Denmark). This is the most common method for collecting small skin samples from free-swimming whales, and has no negative short or long-term effects on individuals other than an initial “startle” response in some cases (Best et al. 2005; Noren and Mocklin 2011; Kowarski et al. 2014). Biopsy samples were collected in association with photo-identification data, as part of ongoing photo-identification studies in all areas. Samples were preserved in either a salt-saturated 20% DMSO solution (Seutin et al. 1991), flash frozen in liquid nitrogen, in RNAlater (Qiagen), or in Allprotect (Qiagen) and frozen upon arrival in the laboratory. All samples were archived in −80°C freezers at the Freshwater Institute (DFO, Canada, Winnipeg) laboratories until further analysis. Bowhead whale biopsy samples were collected under permit Department of Fisheries and Oceans License to Fish for Scientific Purposes S-16/17 1005-NU and Animal Use Protocol FWI-ACC-2016-09.

### 2.2 Genetic Analyses

DNA was extracted using a modified version of the standard Qiagen extraction protocol (Wang et al. 2008). DNA was quantified using a *μ*Quant spectrophotometer (Bio-Tek) or a NanoDrop spectrophotemeter (ThermoFisher Scientific) and normalized to 10–100 ng/*μ*l. Sex was determined through the amplification of a zinc finger gene intron using primers LGL331 and LGL335 (Shaw et al. 2003). PCR fragments were size-separated in 1.5% agarose gels and visualized with GelRed (Biotium). The resulting banding pattern of X (~975 bp) and Y (~1040 bp) fragments were used to infer sex.

Multiplex reactions were developed and used to amplify 23 microsatellite loci (**Table S2**). The reaction mixtures contained 1.0 *μ*l of template DNA, 1X AmpliTaq Gold Buffer, 2.0 mM MgCl_2_, 0.25 mM each dNTP, 0.5 U AmpliTaq Gold DNA polymerase (Life Technologies), and the primer concentrations indicated in **Table S2**. Thermal cycler profiles included a five minute initial denaturing step at 95°C followed by 35 cycles of: 95°C for 30 seconds, annealing temperature (**Table S2**) for 30 seconds, and 72°C for 30 seconds; followed by a final extension step of 72°C for 30 minutes. PCR products were size-separated and visualized on an Applied Biosystems 3130xl Genetic Analyzer (Life Technologies), with GeneScan 600 LIZ (Life Technologies) as a size-standard. Size-standards were checked, and allele calls made using the program GeneMarker (SoftGenetics).

### 2.3 Identifying Recaptures

The crux in identifying genetic recaptures is developing appropriate criteria for defining when two genotypes can be considered the same, and thus representing the same individual. This problem has a long history, and has been most thoroughly dealt with in human forensic DNA typing (National Research Council 1996). The issues are two-fold. First, two genotypes may be the same at the typed loci but represent different individuals if not enough resolution is available with the chosen markers, if individuals are related, or just by chance (a false inclusion). Second, two genotypes may be from the same individual but be scored differently due to genotyping errors or null alleles (a false exclusion). Thus, the key is identifying what criteria minimize the chances of each of these errors.

To identify recaptures, or samples likely originating from the same individual, we used the R package AlleleMatch (Galpern et al. 2012). This package deals with these issues by calculating the genetic similarity of all genotypes, and then creating clusters based on genetic similarity. It then re-creates these clusters by sequentially allowing an incremental number of mismatches, while keeping track of how many “unique” genotypes are present under each number of allowed mismatches. The appropriate number of allowed mismatches is then identified as the point when each genotype associates with only one cluster.

### 2.4 Estimating Abundance

We used Bayesian methods to estimate abundance from the genetic mark-recapture data, based on the general approaches described in Williams et al. (2002) and Kéry and Schaub (2012). We tried two general strategies, each with their own strengths and weaknesses. First, the main goal of this study was to develop a method to use data from sampled locations to explicitly infer the number of individuals that also exist in unsampled locations, and combine these to obtain an estimate for overall abundance. We call this the “location-specific” approach, which was modelled after that used in Durban et al. (2005). The strength of this approach is that it provides an explicit and quantitative means to infer the number of individuals using unsampled locations. The main weakness is that it requires dividing the data by locations, with the consequence that many locations may not have enough samples to be included in the analyses, and thus some information may be lost. The second and most simple approach involved ignoring the location information and treating the data as one large capture-mark-recapture study (hereafter called the “location-independent” approach). The main benefit of this approach is that it allows all samples to be used, regardless of whether recaptures were found within each location. The main weakness of this approach is that it does not explicitly account for the fact that not all locations have been sampled, and therefore makes the assumption that the sampled locations represent adequate sampling opportunities for all individuals. In this case, the location-specific analyses serve as the focus of our work, whereas the location-independent analyses serve as references against which the location-specific estimates can be compared. Both of these methods are described in more detail below.

Again, the main goal of this study was to develop and test an approach that allows for the inference of “missing” individuals when not all sites are sampled. For this approach, we first needed abundance estimates for each sampled location. At this stage, we chose to do this using closed population models for a number of reasons, with the realization that incorporating this approach into open population models is desirable in the future. First, the simplest and clearest way to incorporate our approach, and test its performance, was with closed population models. We therefore felt that this simplest approach was the best way to initialize progress into inferring missing individuals, with the recognition that incorporating it into more complex and realistic models in the future will be required. Second, sampling effort varied widely across locations, in terms of the number of years samples were collected in each location (**Table S1**), as well as the effort put into collecting samples, violating assumptions of open multistate models (Williams et al. 2002). Third, our method accounts for movement among locations, including unsampled ones; given that estimating movement rates (but not explicitly accounting for unsampled locations) is a key aspect of open population models, it is not immediately clear how these could be combined. Therefore, we view this work as a starting point rather than an end point, where future effort can be directed towards incorporating our approach into more complex (and realistic) open population models. Additionally, the long life-span and generally slow life history of bowhead whales mean that it is unlikely that there were dramatic changes to population size (via births, immigration, deaths, or emigration) throughout the study period (Nerini et al. 1984; George et al. 1999; Zeh et al. 2002; George and Bockstoce 2008). However, to try to account for any potential changes over time, for each strategy (location-independent and location-specific) we estimated population sizes based on two data sets: the full data set, and a subset containing only five years of data. The rationale is that the population may have changed throughout the 19 years through which the samples were collected. Therefore, closed-population abundance estimates based on the entire study period may be biased. The 5-year data set contains fewer data but is also expected to be less biased by changes in population size within the sampling period. Combined, this resulted in four estimates of abundance based on: (1) location-specific full data set (LS-FD); (2) location-specific 5-year data set (LS-5Y); (3) location-independent full data set (LI-FD); and (4) location-idependent 5-year data set (LI-5Y).

For all analyses, the recaptures identified by AlleleMatch were filtered to exclude recaptures from the same location in the same year. Bowhead whales are known to vary their migration and habitat use patterns depending on age, sex, and life-history stage (Moore and Reeves 1993; Heide-Jørgensen et al. 2010). Additionally, once in an area, whales are likely to remain there for some period of residency (e.g., Finley 1990). Thus, we thought that within-year recaptures within locations would be artificially high and would not be representative of general capture probabilities for the entire population throughout the study period.

#### 2.4.1 Location-Specific Full Data Set (LS-FD)

Our location-specific approach for estimating abundance was modeled after that used by Durban et al. (2005). The general idea is that location-specific abundance estimates can be used in association with estimated movement rates between locations to identify (or infer) the number of individuals with different sighting histories such as those shown in **Table 1**. When some areas are unsampled, the goal is to estimate the number of individuals in the last row (who were not captured in any area). This abundance estimate can then be used in association with estimates of movement rates to obtain an estimate of total abundance even when some areas remain unsampled. The entire process is summarized in **Fig. 3**, with details described below. There may be more than one unsampled location (i.e., it need not represent a single actual location, but rather could be a number of unsampled locations); however, these will be lumped into a single estimate of “missing” individuals (across all unsampled locations) using this approach. This is because the criterion for being counted as “missing” is that individuals are not present in any of the sampled locations, and therefore the number of unsampled locations need not be defined.

**Figure 3.**
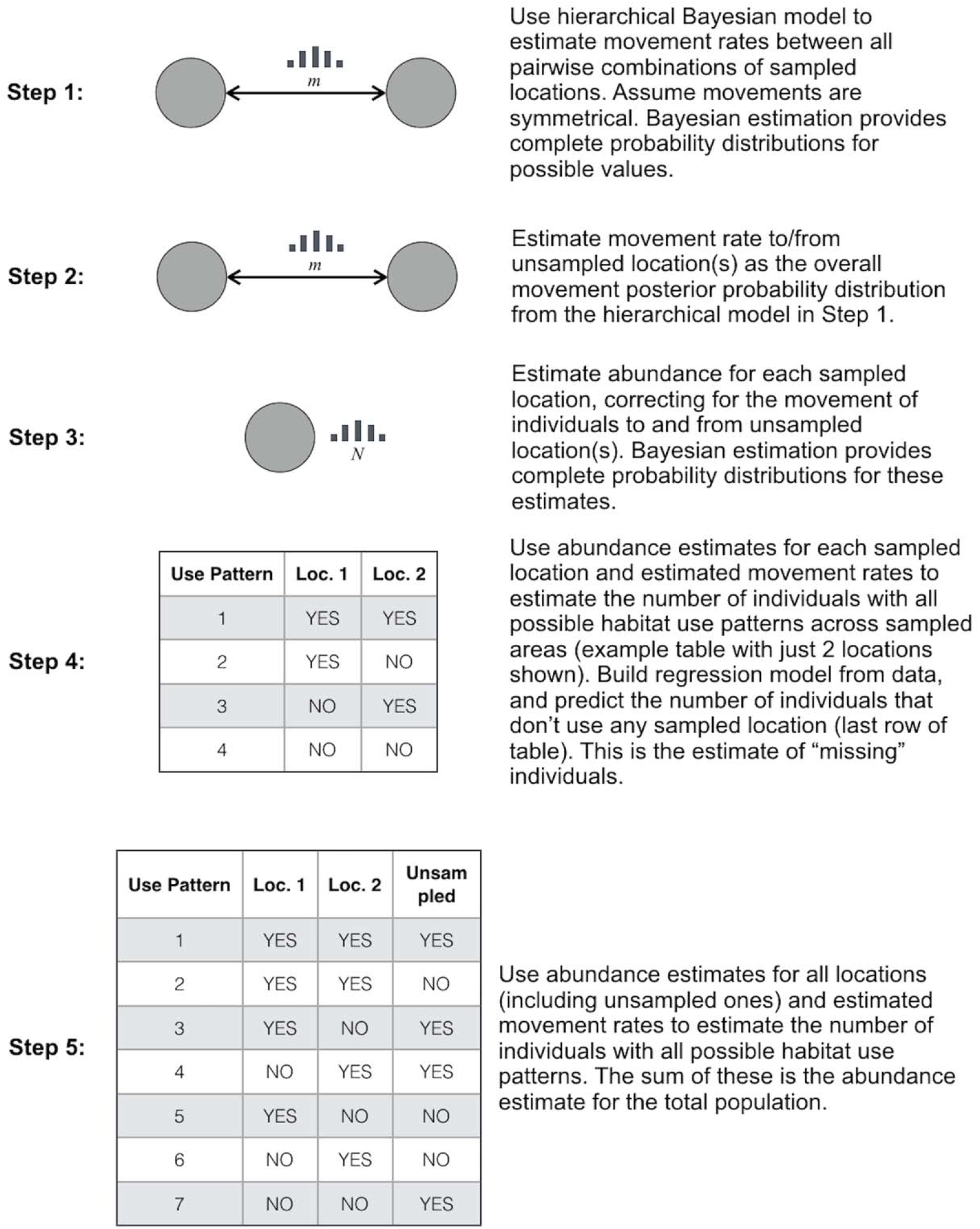
Illustration and brief description of the main steps of the model.

The first step in this process was to estimate the movement rates between all pairs of sampled locations. This is based on the number of recaptures within each considered location, and the number of those recaptures that represent individuals originally marked in the alternative location. We made the assumption that pairwise movement rates were symmetrical, largely because the number of cross-location recaptures was very low (see Results), and therefore there were few data available from which to estimate movement rates. However, it is possible to relax this assumption in the future, which would just require the additional estimation of direction-specific movement rates for each pair of locations. The number of recaptures within and between each pair of locations was converted into a string of zeros and ones, where ones represented recaptures across locations. The posterior probabilities for the movement rates were then estimated using a hierarchical Bayesian approach with a Bernoulli likelihood function and a beta prior with a uniform (flat) distribution (the code itself, as well as a step-by-step walk-through of all analyses are available in the Supplementary Material). The result is a complete posterior probability distribution for movement rates between all pairs of sampled locations, as well as an overall posterior probability distribution for movement rates across all sites.

For the second step of the process, the probability distribution for the symmetrical movement rate to and from the unsampled location(s) was estimated as the overall posterior distribution from the hierarchical model described above across all pairs of locations. This approach makes the implicit assumption that the rate at which individuals move among the sampled locations is at least somewhat representative of the rate at which individuals move between sampled and unsampled locations.

For the third step, abundance was estimated for each sampled location. The genetic recapture data from Allelematch were first converted to typical sighting histories consisting of ones and zeros for each sampling period where individuals were, or were not, captured, respectively. However, these sighting histories are biased due to individuals moving to and from the unsampled location(s). Individuals marked in the sampled location could move to an unsampled location, and therefore not be available for recapture, artificially reducing the number of recaptures. In the opposite direction, unmarked individuals from the unsampled location(s) could move into the sampled location, which would also artificially reduce the proportion of recaptures. Both of these processes reduce the proportion of individuals available for recapture, and would therefore bias population size estimates upwards. To correct for this, for each sampled location we estimated the number of individuals moving to and from the unsampled location(s) and randomly add these to the sighting history data. For example, suppose at one location there were 300 marked individuals and 68 recaptures. Suppose that the estimated movement rate to and from the unsampled location(s) is 0.12. This means that we have “lost” 12% of marked individuals to unsampled locations, and our number of unmarked individuals is also artificially inflated by 12% due to movement of unmarked individuals from unsampled areas into this area. This means that our proportion of recaptures is underestimated by two times the movement rate (24%). Thus, our original 68 recaptures only represent (1 – 0.24 = 0.76) 76% of the recaptures we “should” have had, if individuals did not move to and from unsampled locations (i.e., we should have had 68 / 0.76 = 89 recaptures). To account for this, these (89 – 68 = 21) additional recaptures were randomly added to the sighting histories for this location.

Once the sighting histories were corrected, abundance was estimated for each sampled location using the same closed population approach. Note that although the model assumed a closed population, movement to and from other locations has already been taken into account, and therefore “closed” in this sense just refers to births and deaths. Specifically, to estimate abundance for each location the sighting histories were augmented by 10,000-15,000 (based on trials) to ensure that considered probabilities covered a wide enough range to ensure adequate sampling of the posterior distribution (Tanner and Wong 1987; Royle et al. 2007). This augmenting strategy was introduced by Royle et al. (2007) to deal with the fact that Bayesian estimation of *N* has an unbounded upper end, which can make the efficient exploration of parameter space difficult for many Markov Chain Monte Carlo samplers. Instead, they proposed augmenting data sets by adding a large number of “uncaptured individuals” (individuals with all zeroes in their sighting history), to the observed sighting histories in a manner so that the augmented data set >> *N*. This converts the problem from estimating an unbounded *N* directly into one of estimating what proportion of the augmented data set should be included in the “true” population size (*N*). The parameters of interest are the sighting probability (*p*), and the inclusion probability (Ω), which is the probability that a member of the augmented data set is part of the “true” population size (*N*). Abundance estimates were then obtained for a closed population model (model M_0_ from Otis et al. (1978)) using the approach described in Royle et al. (2007) and Kéry and Schaub (2012). The priors for both the sighting probability (*p*) and the inclusion probability (Ω) were uniform distributions ranging from 0 to 1. All Bayesian analyses were conducted using a combination of R v.3.5.0 (R Core Team 2015), rstan v.2.18.2 (Stan Development Team 2018), and Stan v.2.17.0 (Carpenter et al. 2017). The performance of models for all analyses were assessed based on examination of effective sample sizes, trace plots, and Rhat calculations. Specifically, for all analyses of all scenarios, all Rhat values were 1, all effective sample sizes were greater than 4,000, and all traceplots showed that the chains quickly converged onto one value (and the same value across all chains), and then mildly fluctuated around that value for the vast majority of steps (except the very first few).

For step four, the posterior probabilities for abundance for each sampled location, and for each pairwise movement rate, were used to estimate the number of individuals with each possible habitat use pattern across the sampled locations (there are 3 such possible patterns for 2 locations – **Fig. 3**, 7 possible patterns for 3 locations, and so on). These data were then used in a Bayesian regression analysis to estimate the relevant regression coefficients (see equation below). In this case, *y* is a vector of integers containing the number of individuals with each sighting history, and *x*_1_ through *x*_4_ are vectors of 1s and 0s indicating if each location (locations 1 through 4) is (1) or is not (0) part of each sighting history, respectively. Once estimates for the coefficients (*β_0_* through *β*_4_) have been obtained, these are used to predict the number of individuals that are not seen in any of the sampled locations (i.e., where values for all *x*_1_ though *x*_4_ are zero: the last row of the table shown in **Fig. 3**, step 4). The resulting posterior probability distribution represents the estimate of the total number of individuals not seen in any sampled location.

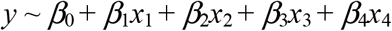

The estimated number of “missing” individuals from the previous step cannot just be added to the estimates for each independent location to obtain an overall abundance estimate. This is because the same individuals may use multiple locations, including a mixture of unsampled and sampled locations (the estimate above is just an estimate of the individuals that *never* use sampled locations, and will therefore often be an underestimate). Therefore, for the fifth and final step, total abundance was estimated in a similar manner as that for the “missing” individuals. Specifically, the complete posterior probabilities for abundance for each location, and for each pairwise movement rate, were used to estimate the number of individuals with each possible habitat use pattern across locations. The difference is that now this table includes data for the unsampled location(s): effectively adding another location to the table and associated movement rates (see **Fig. 3**). The sum of these values then represents the abundance estimate for the total population. Note that one benefit of this approach is that the uncertainty in each estimate is carried over into the uncertainty of the total abundance estimate. Put another way, rather than estimating the total abundance based on the mean, mode, or some other summary metric of the other parameters, this estimate is based on the entire probability distribution for each parameter. In this way the uncertainty in each estimate of abundance and movement are incorporated into the posterior probability distribution for the total abundance estimate. This is one of the attractive properties of setting up the analyses in this way.

#### 2.4.2 Location-Specific 5-Year Data Set (LS-5Y)

We conducted the same analyses as above (for the location-specific full data set, LS-FD), except this time with a data set containing only those sampling events in the 5-year period from 2008–2012. Again, the idea for this process was to reduce the potential biases in the longer-term data set due to births and deaths occurring within the longer time frame, by analyzing data from a much shorter time period (note that immigration and emigration are dealt with explicitly in the model).

#### 2.4.3 Location-Independent Full Data Set (LI-FD)

To estimate abundance for the entire data set, the recapture data from AlleleMatch, combining data across all locations, were first converted to typical sighting histories consisting of ones and zeros for each sampling period (year) where individuals were, or were not, sampled, respectively. These sighting histories were augmented by 30,000 to ensure that considered probabilities covered a wide enough range to ensure adequate sampling of the posterior distribution. An estimate of abundance was then obtained using the same approach as described for the closed population model aspect of the location-specific data set, where estimates of the sighting probability (*p*), inclusion probability (Ω), and abundance (*N*) were obtained.

#### 2.4.4 Location-Independent 5-Year Data Set (LI-5Y)

As with the location-specific 5-year data set, we conducted the same analyses as described above for the location-independent full data set, but this time limiting the data to the years 2008-2013. Again, the rationale was to reduce the potential biases in the longer-term data set changes in population size during the longer sampling time.

### 2.5 Model Validation

A number of simulations were conducted to assess model performance, and to provide information that would be useful for guiding sampling strategies in the field. All simulations were based on the location-specific model, which, instead of estimating abundance for a single population/location, is designed to estimate abundance for each location independently (including unsampled locations), and then come up with one total abundance estimate for all sampled and unsampled locations.

The first set of simulations focused on assessing the effects of varying sampling effort. These simulations were based on four sampled locations and one unsampled location to resemble the available bowhead whale data. For these, all population sizes (including that of the unsampled location) were set to 1,000 individuals (resulting in a total population size of 5,000 individuals). All movement rates between locations were also set to 10%. These initial values were selected to be fairly similar to the expected values for the bowhead whales, and therefore to provide information regarding how this approach would perform with our actual data set.

Additionally, all simulations were run with one capture period and one recapture period, to resemble the bowhead whale data, where no individual was recaptured more than once. Abundance was then estimated based on sampling 5%, 10%, 15%, 20%, and 25% of individuals from each sampled location, to assess how changing sampling effort influences the precision and accuracy of abundance estimates.

A second set of five simulations were also run and analyzed to ensure that the model performs well even when there are large differences in population sizes and sampling effort between locations. As with the first set, these consisted of four sampled locations and one unsampled location. However, this time the abundance at each location (including the unsampled one) was randomly drawn from a list of 20 values ranging from 500 to 10,00, in increments of 500. The movement rates were all set to 10%, and 250 individuals were sampled at each location. This means that the proportion of individuals sampled could range from 2.5% (for a population size of 10,000 individuals) up to 50% (for a population size of 500 individuals).

A third set of two simulations were conducted, but in this case both the population sizes and movement rates were randomly generated. These were conducted to assess how the model performs when there are substantial differences in abundance at different locations, as well as in the movement rates between these locations. Specifically, abundance for each of the four sampled and one unsampled locations were randomly drawn from a list ranging from 250 to 10,000 in increments of 250. Movement rates were similarly randomly drawn from a list ranging from 0.1 to 0.30 in increments of 0.01. The proportion of sampled individuals at each of the sampled locations was set at 0.20.

A fourth set of six simulations was conducted to assess model performance if the movement rate and/or abundance at a single unsampled location differed substantially from those of the sampled locations. For the first scenario the abundance at the four sampled and one unsampled location were all set to 5,000 individuals, but the movement rates among the sampled locations was set to 0.20, and those between the sampled locations and unsampled location were set to 0.05. We then conducted a second simulation reversing this pattern, where the movement rate among the sampled locations was low (0.05), and those between the sampled and unsampled locations was high (0.20). The third simulation consisted of the same movement rates among sampled and unsampled locations (0.15), but all sampled locations had 5,000 individuals, and the unsampled location had just 500. For the fourth simulation this pattern was reversed, with all sampled locations having 500 individuals and the unsampled location having just 500. The movement rate was uniform across all sites at 0.15. For the last pair of simulations under this scenario, both the abundance and movement rates differed between sampled and unsampled locations. Specifically, in the fifth simulation the sampled locations all had 5,000 individuals and the usampled location had 500. The movement rate among the sampled locations was high (0.20), whereas the movement rate between the sampled and unsampled locations was much lower (0.05). For the sixth simulation the population sizes were the same as above, but the movement rates were reversed: the movement rate among sampled locations was 0.05, and that between the sampled locations and the unsampled location were 0.20. Under all six of these scenarios 15% of individuals were sampled from each sampled location.

To gain insight into how the model performs when there is more than one unsampled location, a fifth set of two simulations was conducted and analyzed to assess model performance when there is more than one unsampled location. Specifically, these simulations extended the strategy of randomly generating abundance at each of four sampled locations, as well as at two unsampled locations. The range of values for abundance at each location were randomly drawn from a list ranging from 250 to 10,000 in increments of 250. Movement rates were randomly drawn from a list ranging from 0.05 to 0.30 in increments of 0.01. The proportion of sampled individuals at each of the sampled locations was set at 0.15.

## 3. Results

### 3.1 Samples, Genotypes, and Recaptures

A total of 1,177 samples were genotyped. **Table 2** shows how these were divided among locations. The final microsatellite data set contained 21 loci since two loci were dropped due to a significant portion of the samples missing data that were profiled before 2008. Furthermore, a sample was only included in the analysis if it had at least 10 of the 21 loci scored. This provided an overall probability of identity of 1.16×10^−9^, which corresponds to a 1 in 8.62×10^−8^ chance that the match identified was due to two random animals matching at all loci. **Table S1** shows how these samples were divided by year, location, and sex.

**Table 2.**
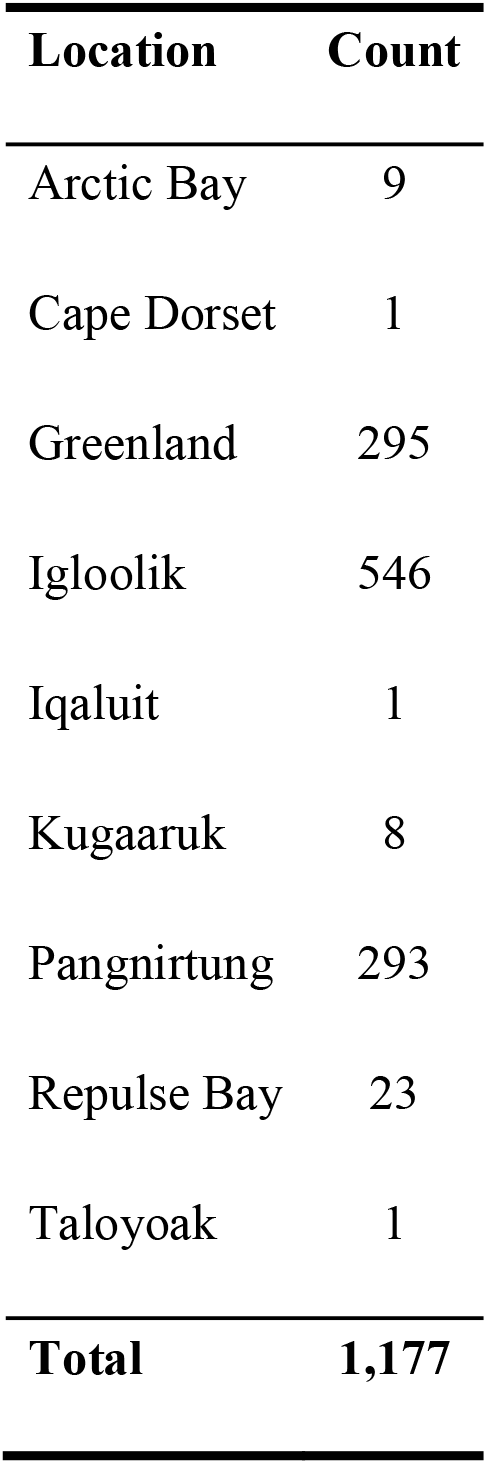
Number of genotyped samples by location.

The AlleleMatch analysis suggested that the best fit for the data was to allow 3 mismatches. Based on this criterion, 992 unique genotypes were identified, along with 185 recaptures. However, 136 of these recaptures represented whales recaptured within the same area and year in which they were marked, and were therefore removed from the analyses. This left 49 recaptures for subsequent analyses. **Table 3** summarizes the recaptures by location. Based on the number of recaptures, only Greenland, Igloolik, Pangnirtung, and Repulse Bay were used for estimating abundance based on the full data set.

**Table 3.** Identified recaptures by location. Only locations with recaptures are included. Location of first capture is indicated by row, whereas location of recapture is indicated by column (i.e,. there where two whale first “captured” in Greenland that were later recaptured in Pangnirtung).

|  | **Greenland** | **Igloolik** | **Pangnirtung** | **Repulse Bay** |
| --- | --- | --- | --- | --- |
| **Greenland** | 5 | 0 | 2 | 0 |
| **Igloolik** | 0 | 23 | 6 | 0 |
| **Pangnirtung** | 0 | 0 | 3 | 3 |
| **Repulse Bay** | 0 | 0 | 0 | 1 |

### 3.2 Model Validation

The results for the simulations testing the effects of sampling effort are shown in **Fig. 4**. The precision of the estimates (or the range between to upper and lower 95% HDI values) declines rapidly until sampling reaches ~15%. Above this sampling effort, precision increases at a much slower rate. Thus, ~15% seems to be a turning point above which diminishing returns are achieved as sampling effort increases. The details of all of the estimates from these simulations are shown in **Fig. 5**. These plots show that the model performs quite well in all aspects (abundance estimation of sampled locations, abundance estimation of unsampled location(s), and total abundance).

**Figure 4.**
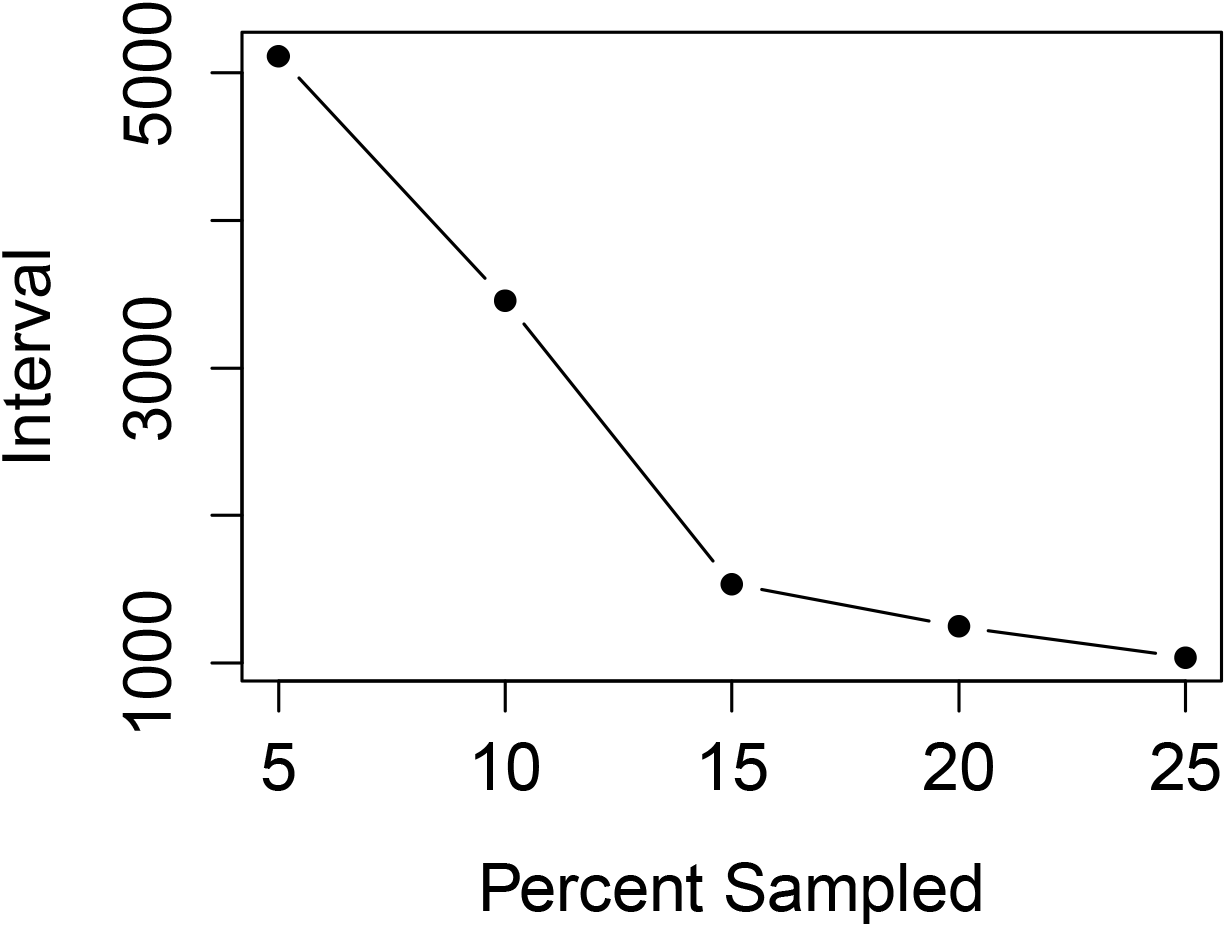
Range of abundance estimates (interval between the upper and lower 95% HDI values) obtained for total abundance when 5, 10, 15, 20, and 25% of the individuals from each location are sampled.

The results from the simulations testing model performance when population size and sampling effort varies across locations are shown in **Fig. 6**. The results show that the model performs quite well under a range of variation in population sizes and sampling efforts. Similarly, the simulations testing model performance when both the abundance and the movement rates varied across all locations showed that the model performs well under these scenarios and recovers accurate estimates of total abundance when there is one (**Fig. S1**), or multiple (**Fig. S2**) unsampled locations.

However, performance declined when the abundance within the unsampled location(s) and/or the movement rates to and from the unsampled location(s) fell well outside the range of values among the sampled locations (**Fig. 7**, **Fig. S3**). This makes intuitive sense because in these cases the information gleaned from the sampled locations (and therefore used to infer information about the unsampled location(s)), is not representative, and therefore leads to inaccurate estimates. For example, when the movement rates to and from the unsampled location were very low (0.05), but those among the sampled locations were high (0.20), the abundance in all locations were biased downwards (**Fig. 7a**). Abundance estimates for the sampled locations were biased downwards because the recapture rates were over-corrected. Specifically, the model assumed a higher movement rate to/from the unsampled location, resulting in an overcorrection of recapture rates for individuals presumed to move to/from the unsampled locations, and therefore resulting in lower abundance estimates. Abundance was underestimated for the unsampled location for the same reason: a higher rate of movement was assumed, resulting in higher estimates of individuals that would be “captured” in the sampled locations, and a lower estimate of those going “missing”. Similarly, when this scenario was reversed and the movement rates to/from the unsampled location were high (0.20), and those among the sampled locations were low (0.05), abundance was overestimated in most locations, and particularly for the total population size (**Fig. 7b**). The rationale is the same as the first scenario, but now recaptures within the sampled locations were under corrected, resulting in overestimates, as well as an underestimation of the probability that individuals would move into sampled areas and be “captured”, resulting in higher estimates of total population size. Similarly, abundance estimates were biased if the actual abundance in the unsampled location(s) were vastly outside of the range of those within the sampled locations (**Fig. 7c-d, Fig. S3**). Again, this is expected because the sampled locations are not providing appropriate information on which inferences about the unsampled locations can be made.

### 3.3 Bowhead Whale Abundance Estimates

The location-specific abundance estimates for each location, inferred for the unsampled location(s), and for the total population, are shown in **Table 4** (rows 1–6). The mode was used as the point estimate instead of the mean in all cases because the posterior probability distributions for all abundance estimates were skewed, with heavy right-hand tails, and therefore the mode represented the estimate with a higher probability than the mean. Note that the estimates for each location are inclusive, meaning that they represent all whales that use each area (and may also be counted for other areas), rather than representing whales that only use each area. The locationin–dependent abundance estimates based are also shown in **Table 4** (row 7).

**Table 4.** Abundance estimates (modes of the posterior probabilities) for each location, as well as for the full population. Included are these estimates and 95% highest density intervals (HDI). Total-LS refers to the abundance estimate for the total population based on inference from the location-specific analyses and Total-LI refers to estimates based on the location-independent analyses.

| Row # | Location | Full Data Set |  | 5 Years of Data |  |
| --- | --- | --- | --- | --- | --- |
|  |  | Estimate | 95% HDI | Estimate | 95% HDI |
| 1 | Greenland | 2,615 | 1,353–6,114 | 2,089 | 928–5,748 |
| 2 | Igloolik | 4,201 | 2,952–6,252 | 2,571 | 1,698–4,024 |
| 3 | Pangnirtung | 2,378 | 1,491–4,203 | 2,852 | 1,210–7,868 |
| 4 | Repulse Bay | 35 | 20–90 | NA | NA |
| 5 | “Missing” | 449 | 0–8,297 | 4,111 | 179–19,721 |
| 6 | Total-LS | 11,747 | 8,169–20,043 | 13,899 | 7,782–30,602 |
| 7 | Total-LI | 11,682 | 8,620–16,014 | 6,877 | 4,828–11,477 |

## 4. Discussion

Analyses of simulated data sets suggest that our new approach performs quite well, and could be a useful tool for estimating abundance of other populations where only a subset of possible locations have been sampled. Simulations testing analysis performance with respect to sampling effort show that sampling roughly 15% of the individuals at each location is desirable, whereas sampling more intensively will result in diminishing returns. The approach still performs well when fewer than 15% of individuals are sampled at each location, but there is decreasing precision around the estimates as sample size decreases (**Fig. 5**).

**Figure 5.**
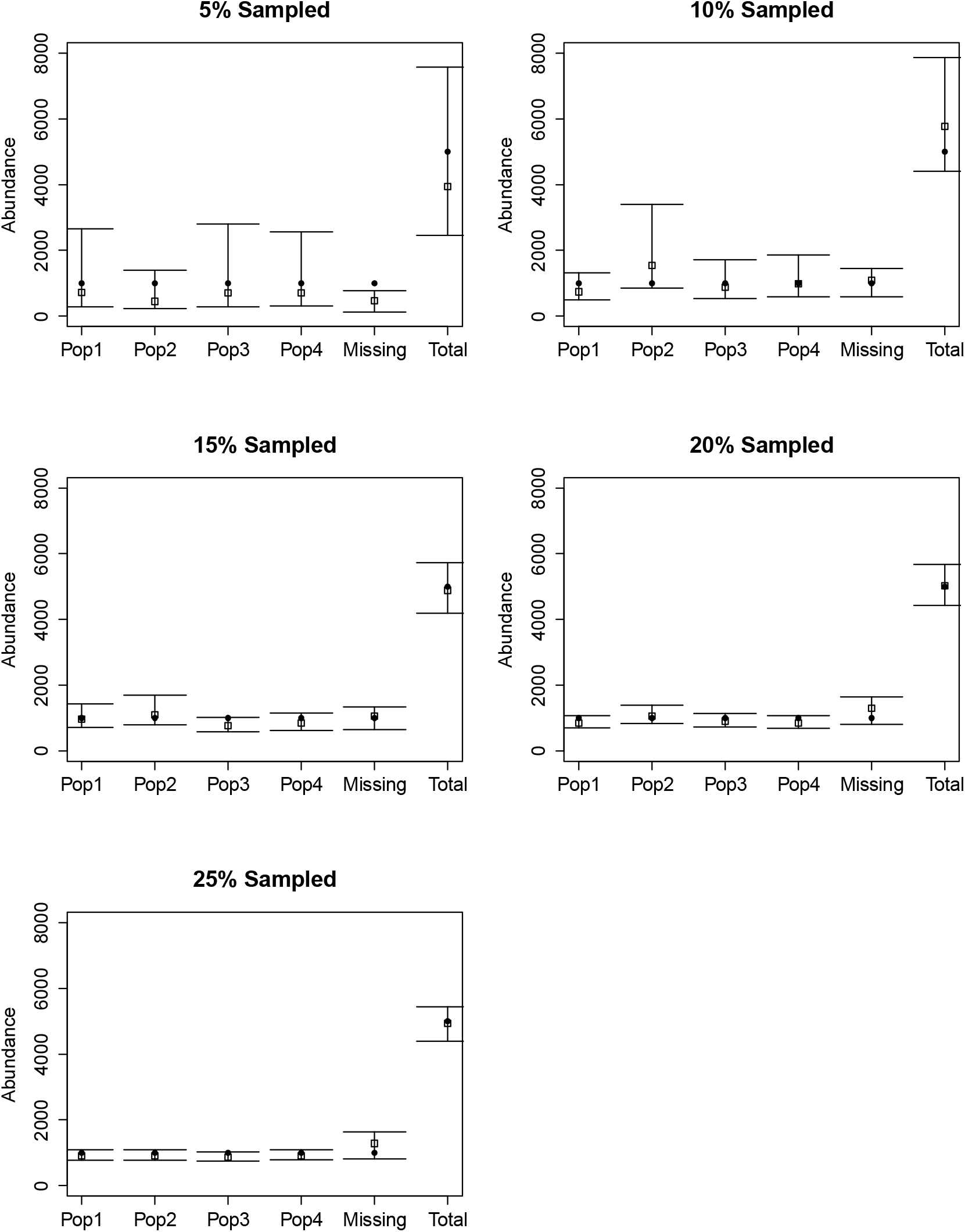
Abundance estimates and associated 95% HDIs for each simulated scenario testing the effects of sampling effort. Actual values are shown in black circles. Here “Missing” refers to the number of individuals in the unsampled location(s).

The approach also performs well even when there is wide variation in abundance between sampling locations. Although the simulations used in our tests were not exhaustive, they do show that accurate abundance estimates can be obtained for the total population when only a subset of locations are sampled, and when there is wide variation in abundance across locations (**Fig. 6**). Combined, the analysis of simulated scenarios suggests that this new approach may indeed be a viable option for estimating abundance from genetic mark-recapture data when not all locations are sampled.

**Figure 6.**
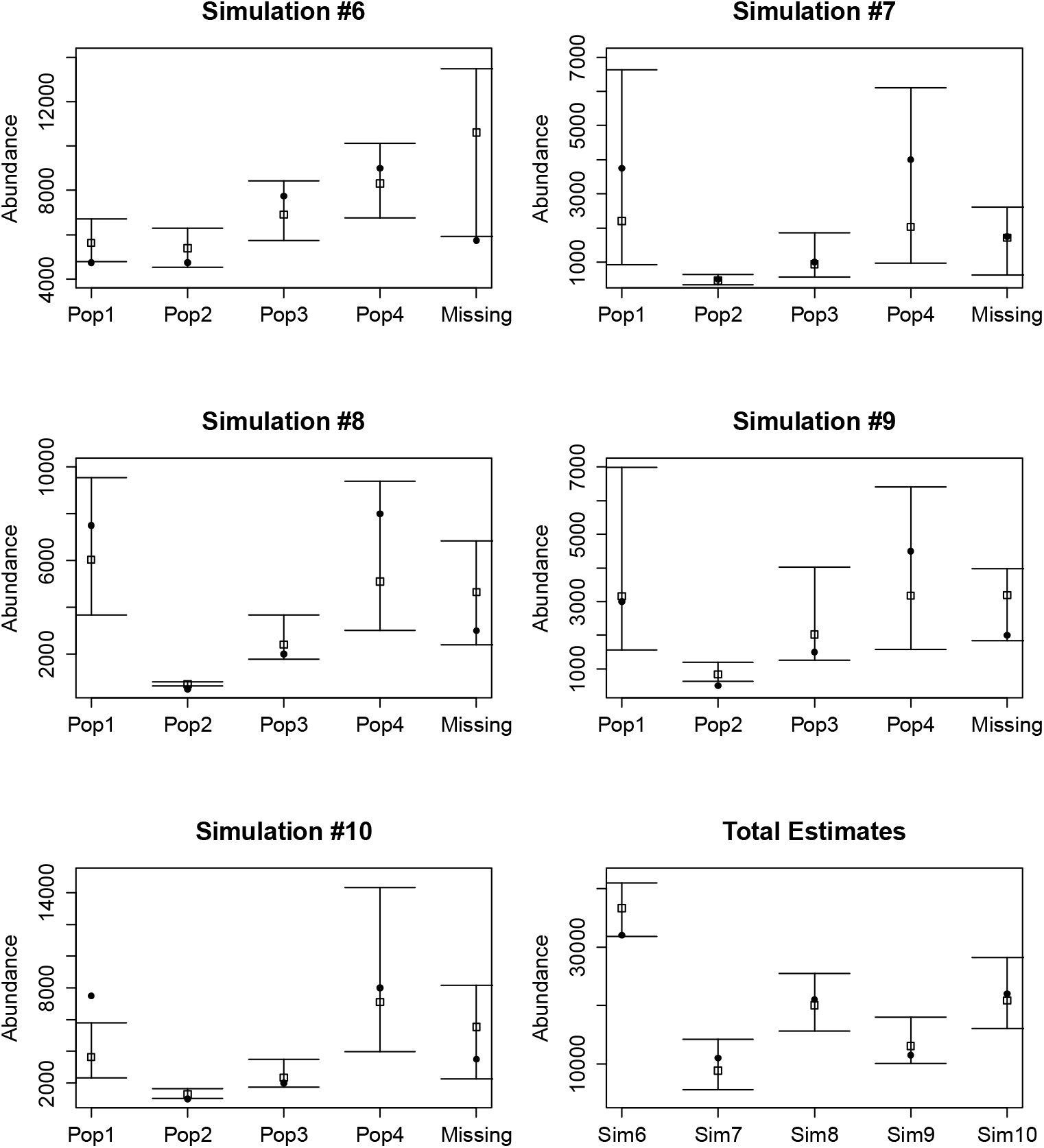
Abundance estimates and associated 95% HDIs for each simulated scenario testing the effects of varying population sizes and sampling efforts across locations. Actual values are shown in black circles. Here “Missing” refers to the number of individuals in the unsampled location(s). Note that the “total” abundance estimate for all 5 tested scenarios are located in the bottom-right panel. These were separated from the other results for each simulated condition because the scale is so different for the total estimates versus those for each location independently.

**Figure 7.**
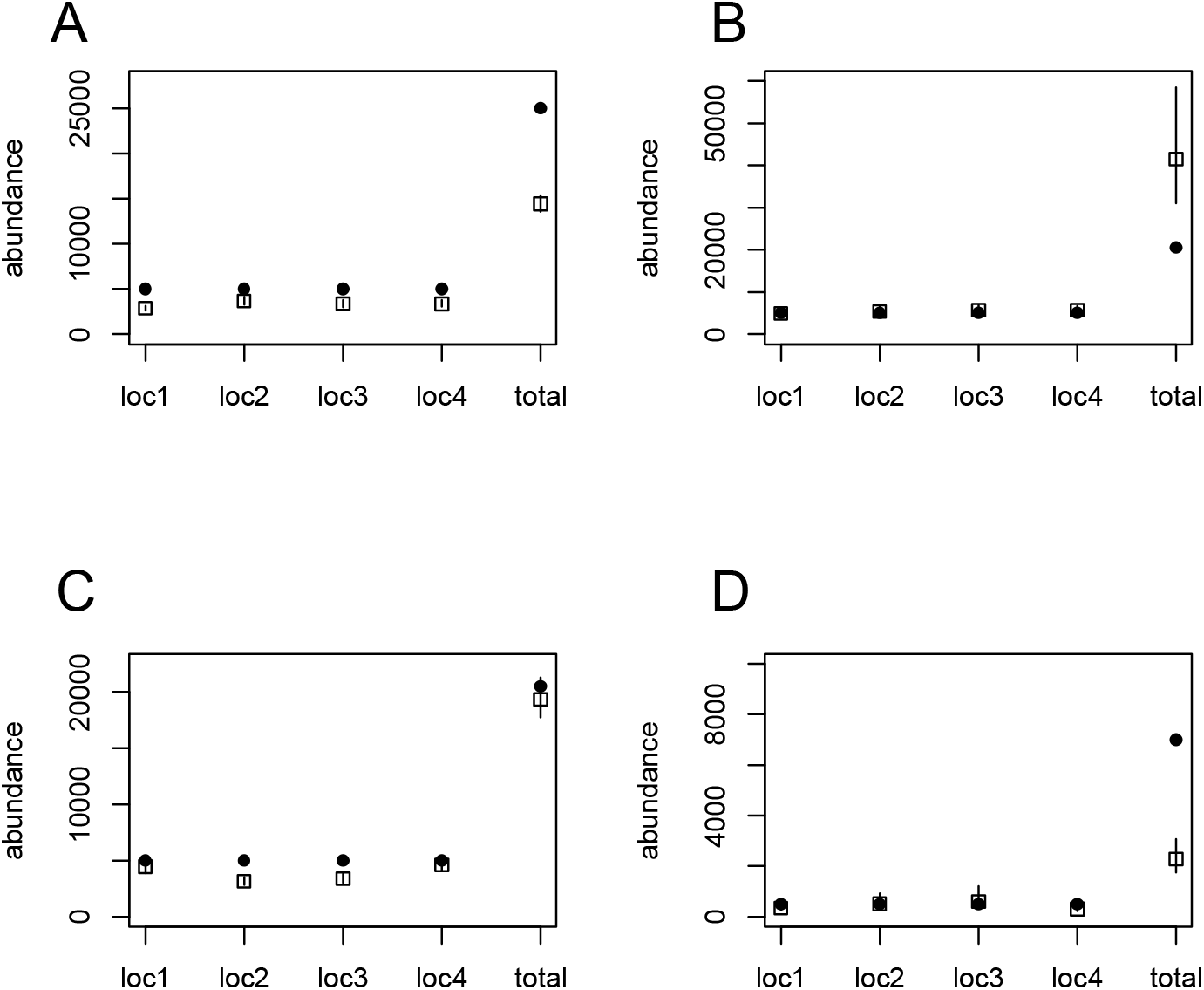
Actual (black circles), and estimated (open squares) abundance for four sampled locations, and total abundance, when there is one unsampled location. Bars represent the 95% HDIs. A. Abundance for each location (4 sampled and 1 unsampled) are 5,000 individuals, for a total of 25,000. The movement rates among sampled locations was high (0.2), and those between the sampled and unsampled locations was low (0.05). B. The reverse situation from A: abundance at each location as 5,000 (for a total to 25,000), but movement rates among sampled locations was low (0.05) and those between sampled and unsampled locations was high (0.20). C. All movement rates were equal at 0.15, but abundance was 5,000 individuals for all sampled locations and 500 for the unsampled location, for a total abundance of 20,500 individuals. D. The reverse of C: all movement rates were equal at 0.15, but abundance was 500 individuals for all sampled locations and 5,000 for the unsampled location, for a total abundance of 7,000 individuals.

Performance of our approach declined, however, when the data from the sampled locations were not representative of those for the unsampled locations. Specifically, if the abundance in the unsampled locations and/or the movement rates to and from the unsampled locations are well outside the range of those among the sampled locations, then our model was unable to obtain accurate estimates. This is due to the fact that the information obtained from the sampled locations is not representative of that in the unsampled locations, which leads to incorrect inferences. Therefore, using this approach carries the implicit assumption that the abundance of, and movement rates to and from, unsampled locations are at least somewhat within the range of those among the sampled locations. Although this seems like a major drawback, it suggests that our method may be a useful approach for studies when abundance estimates are desired, but it has not yet been possible to sample from all areas. Using our approach will produce more accurate abundance estimates than ignoring unsampled sites altogether, but they will obviously not be as accurate as those possible if effort could be distributed across all sites.

For analyses of the bowhead whale data specifically, the number of recaptures was very low, despite the substantial effort in sample collection across a wide range of areas. This fact alone suggests that the population size is quite large, and demonstrates the inherent difficulties in estimating abundance for EC-WG bowhead whales. Given their movement dynamics and heterogeneity, it seems that any estimate of population size is going to have its shortcomings, and have to be taken with a fairly large degree of caution. However, here we have tried to be comprehensive in our analyses to try to make the most of the data that are available.

For the location-specific estimates for sampled locations, our results are comparable to previously published estimates that are available. First, Cosens and Innes (2000) report abundance estimates from aerial line-transect surveys for a larger area centred around Repulse Bay. Their estimate was 75 individuals, with a 95% confidence interval of 17–133. This agrees well with our estimate of 35 (95% HDI 20–90), particularly when considering that they covered a larger area than was available here. Second, Rekdal et al. (2014) report aerial survey line-transect and genetic mark-recapture abundance estimates for West Greenland. The estimates from these methods were 744 whales (95% CI: 357–1,461) and 1,538 whales (827–2,249), respectively. However, the aerial surveys were conducted in a single year, and therefore the resulting estimates do not include whales that may use the area, but did not do so in the year of the survey, which may represent a substantial number. Our genetic mark-recapture abundance estimate for West Greenland is a bit larger (2,615, 95% HDI: 1,353–6,114), and with a heavier right-handed tail. This makes sense given that our approach attempts to infer whales that may be missing, and therefore results in higher probabilities associated with larger estimates. Given the heterogeneity in bowhead movement patterns, we think that basing estimates on a longer-term data set increases the chances of including all whales that use specific areas, and will therefore result in more accurate estimates.

For total abundance estimates, our values were fairly similar across approaches (location-specific and location-independent), as well as across time periods used (full data set versus 5 years of data) (**Table 4**). In most cases the estimates for the 5-year data set were slightly lower than those for the whole data set. This could result from two primary reasons. First, the higher estimates may be correct, but only obtained with the larger data set due to the much larger sample size available. Second, the higher estimates may be biased by population growth during the study period, which would result in abundance estimates for the longer time period being biased upwards. Precision was higher for the longer time period, which was expected due to the larger number of recaptures available for the longer-term data set. Given these differences and potential issues, we think that the location-specific estimate based on the full data set currently represents the best overall estimate from these analyses (11,747 whales, 95% HDI: 8,169–20,043). The precision of this estimate is quite low, indicating that effort should be improved in the future to obtain more samples from these locations. Interestingly, Fisheries and Oceans Canada (DFO) recently (in 2013) carried out large-scale aerial surveys of the arctic to estimate abundance of narwhal and bowhead whale populations. The estimate they obtained for the EC-WG bowhead population was 6,743 individuals (CV = 22%) (Doniol-Valcroze et al. 2014). However, this is likely an underestimate given that several key areas were not covered by those surveys (Foxe Basin, and northern Hudson Bay). Thus, we would expect our estimate to be larger than that reported based on the aerial surveys.

These analyses show that, at least in the tested conditions, it is possible to use regression techniques to infer “missing” individuals in mark-recapture abundance analyses, and therefore improve abundance estimates for situations where it is not possible to sample all locations. This advancement should be useful to a wide range of studies. We recognize that several improvements could be made to this approach, such as incorporating it into open population models, as opposed to just the closed population models considered here. However, we think that it represents an important step forward – and a useful tool – in the quest to obtain accurate estimates of abundance for populations that are difficult to study, but it is not meant to represent a final solution. For studies and management of the EC-WG bowhead whale population, the benefits of this approach may take many other forms as well. For example, the systematic collection and analysis of biopsy samples from a few key locations is vastly less expensive than conducting exhaustive aerial surveys of the high arctic. Thus, this may represent a more efficient means to assess and monitor this population over time than the methods that are currently employed.

## Acknowledgements

Over the years numerous people have been involved in sample collection. This work would not have been possible without their participation. We sincerely thank the Hunters and Trappers Organizations and the communities of Igloolik, Pangnirtung, Arctic Bay, Repulse Bay, Kuugaaruk, Taloyoak and Cape Dorset who provided logistic support, field assistance and/or participation for the biopsy sampling and collection programs. We appreciate the work done by Denise Tenkula (DFO) for laboratory analyses and scoring of alleles. We are grateful to DFO staff for their efforts in receiving, handling and curating of samples. Funding for the sample collection and genetic analyses has been obtained from DFO, Polar Continental Shelf Project, Nunavut Implementation Fund, the Nunavut Wildlife Research Trust, ArcticNet Centre of Excellence, International Polar Year (Global Warming and Arctic Marine Mammals), Assiniboine Park Zoo, the University of Manitoba and Natural Sciences and Engineering Research Council. This work was funded by Fisheries and Oceans Canada (DFO). To comments of two anonymous reviewers greatly improved this manuscript and the degree to which this approach was tested.

## Data Accessibility

The data, code, and instructions for combining the two to obtain the results are available at https://github.com/timothyfrasier/bowhead_abundance. Specifically, there is a pdf file containing “walk-through” instructions, explaining how to conduct the analyses and obtain the results presented in the paper. There are also two folders: one containing the required data, and another containing the required code. Users should be able to replicate our results by following the instructions.

