## Supplementary material for "Abundance estimation from genetic mark-recapture data when not all sites are sampled: an example with the bowhead whale": Fig. S2

**Figure S2.** Results of simulations where abundance for each location (four sampled and two unsampled), and movement rates among all locations, were randomly selected as described in the Methods section. True values are shown in black circles, the mode of estimates are in open squares, and the 95% highest density interval (HDI) of the estimates are represented by lines. **A**. Actual abundance values were 250, 8500, 6250, 5250, and 32750 for the four sampled locations and the total population (including the two unsampled locations), respectively. Movement rates among locations 1-6, where 1-4 represent the sampled locations, and locations 5 and 6 represent the unsampled locations, were: m12 = 0.14, m13 = 0.09, m14 = 0.07, m15 = 0.11, m16 = 0.05, m23 = 0.24, m24 = 0.18, m25 = 0.06, m26 = 0.17, m34 = 0.16, m35 = 0.10, m36 = 0.12, m45 = 0.08, m46 = 0.21, m56 = 0.13. **B.** Actual abundance values were 7500, 5500, 2250, 9750, and 37000 for the four sampled locations and the total population (including the two unsampled locations), respectively. Movement rates among locations were: m12 = 0.24, m13 = 0.07, m14 = 0.09, m15 = 0.23, m16 = 0.25, m23 = 0.08, m24 = 0.19, m25 = 0.21, m26 = 0.12, m34 = 0.18, m35 = 0.06, m36 = 0.16, m45 = 0.15, m46 = 0.14, m56 = 0.22.

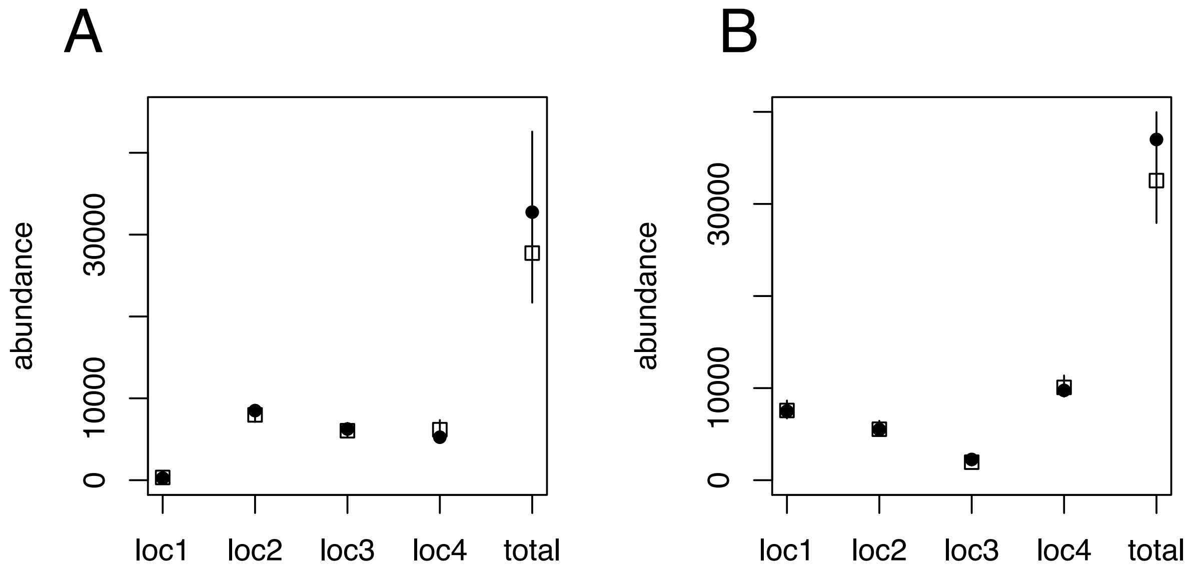
