## Supplementary material for "Abundance estimation from genetic mark-recapture data when not all sites are sampled: an example with the bowhead whale": Fig. S3

**Figure S2.** Results of simulations where both the abundance and the movement rates differ markedly between the sampled locations and a single unsampled location. True values are shown in black circles, the mode of estimates are in open squares, and the 95% highest density interval (HDI) of the estimates are represented by lines. **A**. Actual abundance values were 5000 for all sampled locations and 500 for the unsampled location; movement rates were high (0.2) among the sampled locations but low (0.05) between the sampled and unsampled locations. **B.** , Actual abundance values were 5000 for all sampled locations and 500 for the unsampled location; movement rates were low (0.05) among the sampled locations but high (0.20) between the sampled and unsampled locations.

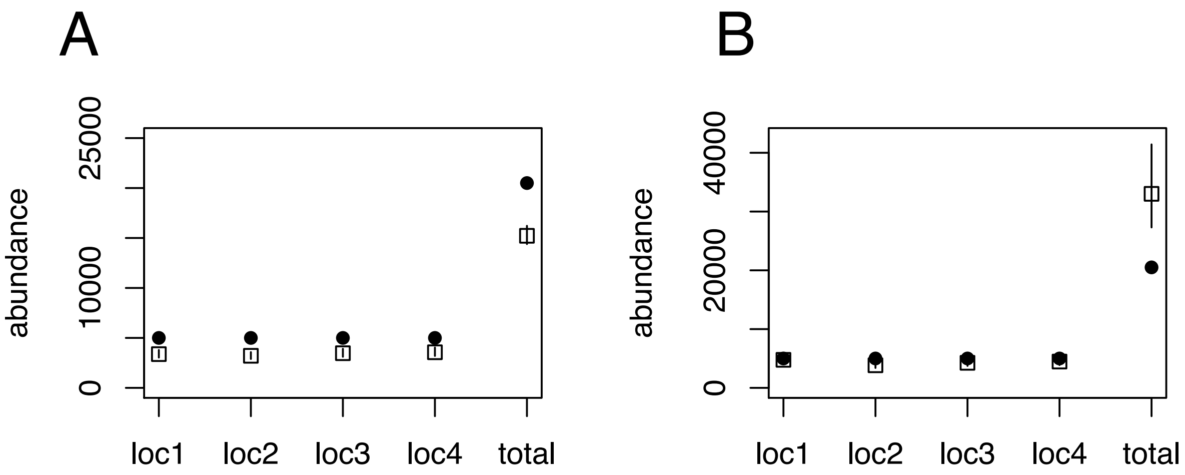
