## Supplementary material for "Abundance estimation from genetic mark-recapture data when not all sites are sampled: an example with the bowhead whale": Table S1

**Table S1:** Sample numbers by year, location and sex.

| Year | Location | Sex | Count |
|---|---|---|---|
| 1995 | Igloolik | Unknown | 10 |
| 1996 | Igloolik | Unknown | 17 |
| 1997 | Igloolik | Unknown | 1 |
| 1997 | Pangnirtung | Unknown | 17 |
| 1997 | Repulse Bay | Unknown | 5 |
| 1998 | Repulse Bay | Unknown | 3 |
| 2000 | Repulse Bay | Unknown | 4 |
| 2001 | Geenland | Unknown | 13 |
| 2001 | Igloolik | Unknown | 34 |
| 2001 | Kugaaruk | Unknown | 2 |
| 2001 | Repulse Bay | Unknown | 3 |
| 2002 | Greenland | Unknown | 13 |
| 2002 | Igloolik | Unknown | 53 |
| 2002 | Kugaaruk | Unknown | 5 |
| 2002 | Pangnirtung | Unknown | 10 |
| 2003 | Greenland | Unknown | 10 |
| 2003 | Igloolik | Unknown | 27 |
| 2004 | Pangnirtung | Unknown | 6 |
| 2005 | Greenland | Unknown | 17 |
| 2005 | Pangnirtung | Unknown | 15 |
| 2005 | Repulse Bay | Female | 1 |
| 2006 | Greenland | Unknown | 22 |
| 2006 | Pangnirtung | Unknown | 31 |
| 2007 | Greenland | Female | 130 |
| 2007 | Greenland | Male | 26 |
| 2008 | Arctic Bay | Female | 2 |
| 2008 | Greenland | Female | 45 |
| 2008 | Greenland | Male | 8 |
| 2008 | Greenland | Unknown | 8 |
| 2008 | Igloolik | Female | 4 |
| 2008 | Igloolik | Male | 2 |
| 2008 | Pangnirtung | Female | 1 |
| 2008 | Pangnirtung | Male | 1 |

| | | | |
|---|---|---|---|
| 2008 | Pangnirtung | Unknown | 1 |
| 2008 | Repulse Bay | Female | 1 |
| 2008 | Repulse Bay | Unknown | 4 |
| 2009 | Arctic Bay | Female | 5 |
| 2009 | Arctic Bay | Male | 1 |
| 2009 | Cape Dorset | Male | 1 |
| 2009 | Igloolik | Female | 42 |
| 2009 | Igloolik | Male | 40 |
| 2009 | Repulse Bay | Female | 1 |
| 2009 | Repulse Bay | Male | 1 |
| 2011 | Igloolik | Female | 17 |
| 2011 | Igloolik | Male | 24 |
| 2011 | Iqaluit | Male | 1 |
| 2011 | Kugaaruk | Female | 1 |
| 2011 | Pangnirtung | Female | 28 |
| 2011 | Pangnirtung | Male | 25 |
| 2012 | Arctic Bay | Male | 1 |
| 2012 | Igloolik | Female | 58 |
| 2012 | Igloolik | Male | 53 |
| 2012 | Igloolik | Unknown | 4 |
| 2012 | Pangnirtung | Female | 38 |
| 2012 | Pangnirtung | Male | 66 |
| 2012 | Pangnirtung | Unknown | 5 |
| 2012 | Taloyoak | Female | 1 |
| 2013 | Igloolik | Unknown | 160 |
| 2013 | Pangnirtung | Female | 14 |
| 2013 | Pangnirtung | Male | 34 |
| 2013 | Pangnirtung | Unknown | 1 |
| Unknown | Greenland | Unknown | 6 |
