## Supplementary material for "Abundance estimation from genetic mark-recapture data when not all sites are sampled: an example with the bowhead whale": Table S2

**Table S2:** Multiplex reactions for the amplification of microsatellite loci. Included is the name, reaction number, primer concentration, annealing temperature, and reference for each locus.

| **Name** | **Reaction** | **Primer [ ]** | $\mathbf{T}_a$ | **Reference** |
|---|---|---|---|---|
| Bmy1 | 1 | 1.6 $\mu$M | 55°C | Huebinger *et al* (2008) |
| Bmy8 | 1 | 0.2 $\mu$M | 55°C | Huebinger *et al* (2008) |
| Bmy16 | 1 | 0.6 $\mu$M | 55°C | Huebinger *et al* (2008) |
| EV37 | 1 | 0.16 $\mu$M | 55°C | Valsecchi & Amos (1996) |
| EV104 | 1 | 0.4 $\mu$M | 55°C | Valsecchi & Amos (1996) |
| Bmy10 | 2 | 0.4 $\mu$M | 55°C | Huebinger *et al* (2008) |
| Bmy55 | 2 | 0.6 $\mu$M | 55°C | Huebinger *et al* (2008) |
| EV76 | 2 | 0.2 $\mu$M | 55°C | Valsecchi & Amos (1996) |
| RW31 | 2 | 1.2 $\mu$M | 55°C | Waldick *et al* (1999) |
| FCB4 | 2 | 0.3 $\mu$M | 55°C | Buchanan *et al* (1996) |
| Bmy19 | 3 | 0.3 $\mu$M | 55°C | Huebinger *et al* (2008) |
| Bmy33 | 3 | 0.4 $\mu$M | 55°C | Huebinger *et al* (2008) |
| Bmy36 | 3 | 0.6 $\mu$M | 55°C | Huebinger *et al* (2008) |
| Bmy53 | 3 | 1.6 $\mu$M | 55°C | Huebinger *et al* (2008) |
| Bmy54 | 3 | 0.4 $\mu$M | 55°C | Huebinger *et al* (2008) |
| Bmy49 | 4 | 0.8 $\mu$M | 58°C | Huebinger *et al* (2008) |
| Bmy58 | 4 | 1.0 $\mu$M | 58°C | Huebinger *et al* (2008) |
| RW18 | 4 | 0.3 $\mu$M | 58°C | Waldick *et al* (1999) |
| Bmy11 | 5 | 0.3 $\mu$M | 58°C | Huebinger *et al* (2008) |
| Bmy57 | 5 | 0.12 $\mu$M | 58°C | Huebinger *et al* (2008) |
| EV1 | 5 | 1.6 $\mu$M | 58°C | Valsecchi & Amos (1996) |
| Bmy12 | 6 | 0.5 $\mu$M | 58°C | Huebinger *et al* (2008) |
| Bmy26 | 7 | 0.5 $\mu$M | 55°C | Huebinger *et al* (2008) |
| GATA098 | 8 | 0.5 $\mu$M | 48°C | Palsbøll *et al* (1997) |
